# The effect of indoor daylight spectrum and intensity on viability of indoor pathogens on different surface materials

**DOI:** 10.1101/2022.01.14.476401

**Authors:** Man In Lam, Kinga Vojnits, Michael Zhao, Piers MacNaughton, Sepideh Pakpour

**Affiliations:** Faculty of Applied Science, School of Engineering, University of British Columbia, Kelowna, BC, Canada; Department of Environmental Health, Harvard T.H. Chan School of Public Health, Boston, MA, USA

**Keywords:** Healthy Built Environment, Indoor Microbiome, Indoor daylight, Smart Window, Pathogens on Surfaces, Pathogens Viability

## Abstract

Built environments play a key role in the transmission of infectious diseases. Ventilation rates, air temperature and humidity affect airborne transmission while cleaning protocols, material properties and light exposure can influence viability of pathogens on surfaces. We investigated how indoor daylight intensity and spectrum through electrochromic (EC) windows can impact the growth rate and viability of indoor pathogens on different surface materials (polyvinyl chloride (PVC) fabric, polystyrene (PS), and glass) compared to traditional blinds. Our results showed that tinted EC windows let in higher energy, shorter wavelength daylight than those with clear window and blind. The growth rates of pathogenic bacteria and fungi were significantly lower in spaces with EC windows compared to blinds: nearly 100% growth rate reduction was observed when EC windows were in their clear state followed by 41-100% reduction in bacterial growth rate and 26-42% reduction in fungal growth rate when EC windows were in their darkest tint. Moreover, bacterial viabilities were significantly lower on PVC fabric when they were exposed to indoor light at EC-tinted window. These findings are deemed fundamental to the design of healthy modern buildings, especially those that encompass sick and vulnerable individuals.

**Practical Implications:**

- Light is an important factor that influences occupant health.
- Healthcare Associated Infections (HAI) bring substantial costs on the healthcare systems hence new disinfection methods are always needed to minimize fomites especially with the increasing antibiotic resistance.
- We found that indoor light modulated by the EC smart windows can significantly reduce the growth rate and viability of pathogenic bacteria and fungi, which is mainly due to the high energy blue light spectrum at wavelength of 400-500nm.
- Pathogenic fungi are found to be more affected by the indoor light intensity, while indoor bacteria on surfaces are more susceptible to the light spectrums.
- These results also demonstrate the promising potential of indoor daylight exposure as an alternative for fomite disinfection strategy and expand the benefits of EC window as part of healthy building design in the future.

## 1 INTRODUCTION

The built environment plays an important role in occupant health as people today spend the majority of their time indoors. Researchers and engineers have been cooperating together over the years to establish a more resource-efficient and healthier built environment for occupants^1^. However, assessing indoor environment quality is complex as buildings serve a variety of functions and many factors need to be taken into consideration. The Harvard T.H Chan School of Public Health established a Healthy Building framework which includes nine foundations: ventilation, air quality, thermal health, water quality, moisture, dust and pest, noise, safety, and lighting and views^1^. These factors act individually and interactively in shaping the indoor environment and affecting the physiological and psychological health of building occupants.

Among the nine foundations of healthy buildings, light is an important attribute affecting occupant’s health and productivity. Namely, light catalyzes hormone secretions in human body, controls our circadian rhymes and thus regulates our sleep, mood, and work performance^2,3^. Therefore, daylight can not only affect thermal and visual comfort levels of the occupants but also influence their cognitive functions and mental health^4^. Inadequate daylight exposure have been frequently associated with poor sleep quality, decrease in productivity, and more workplace errors^5–7^. Windows, which modulate outdoor solar radiation transmission and indoor daylight spectrum, play a key role in determining the build environment quality. Modern buildings often designed to maximize glazing to increase the views, but in practice occupants lower the blinds to control for glare and thermal discomfort^3^. On average, 59% of window area is obstructed by blinds^8^. Moreover, traditional low emission (low-E) glass has a solar heat gain coefficient of 0.48, which means approximately 48% of heat are transmitted into the building^9,10^. Building systems need to be sized to accommodate the hottest summer day, resulting in oversized systems to meet the peak load from solar heat gain in the building^2,3,11^. Studies have shown that up to 60% of energy is lost through the windows, and a range of 10-25% heat loss in the residential buildings are due to the windows^2,12^. Electrochromic (EC) windows have been developed to overcome these issues by embedding EC materials within the window layers and employing multiple-pane glazing^11,12^. When a low voltage electric current is applied to the EC materials, the consequential redox reaction changes the light transmission hence dynamically controlling the indoor light spectrum^11^. EC windows provide similar light transmission to low-E glass when the sun is not on the facade, while tinting to mitigate glare and solar heat gain when the sun is on the facade, which is more sustainable and provides a better occupant experience^7,9,13,14^.

In addition, indoor microbiota which consists of ever-changing combinations of bacteria, virus, and fungi have important relevance to the built environment quality^24^. Although exposure to beneficial microbes in indoor spaces can positively effect occupants’ health, albeit still uncertain on its causality, exposure to indoor infectious agents, via aerosol droplets, direct contact with pathogens, or indirect contact with fomites^25–28^, are known to negatively impact occupants’ health. Contaminated surfaces can serve as reservoirs for pathogens, particularly problematic in hospital settings, and facilitate disease tranmissions^30,31^. Nosocomial infections, also known as the healthcare-associated infections (HAI), are diseases that do not present in patients during the time of admission but acquired during their hospital stays. Although all pathogens are problematic to hospitalized patients, HAIs are often related to only a few bacterial species, including *Enterococcus faecium, Staphylococcus aureus, Klebsiella pneumoniae, Acinetobacter baumannii, Pseudomonas aeruginosa, and Enterobacter faecium*, namely the ESKAPE pathogens^32,33^. Most of the ESKAPE species are on the list of the most problematic microbial species by World Health Organization (WHO) due to their multiple antibiotic resistances^30,32^. This could be accounted by the pathogens’ biofilm-forming ability, which provides the mechanical and biochemical protections for microbes after attaching to inanimate surfaces and make hospital disinfection procedures more challenging^30^. The European Centre of Disease and Control (ECDC) estimated that an additional 900 million Euros hospital costs in European Union in 2007 was due to five of the ESKAPE pathogens, bringing significant financial burden to the healthcare system^32^. Extensive studies have investigated the inactivation methods of EKSPAE pathogens, though nosocomial infections are not only caused by these five bacterial species. Other biofilm-forming species such as *Escherichia coli* and *Staphylococcus epidermis* could also cause nosocomial infections and result in serious health outcomes such as sepsis when treatments are not received on time^32,33^. Fungi, viruses and parasites can also cause HAIs, with immunocompromised patients being the most vulnerable populations^30^. In fact, cases of fungal nosocomial infections have been noticeably increased over the past few decades due to aging populations in developed countries, with more immunocompromised patients being seen and more immunosuppressive agents being used in the hospitals^34,35^. The most common cause of fungal nosocomial infections are *Candida and Aspergillus species*, leading to candidemia and aspergillosis, respectively, both with high mortality rates^34^. Unfortunately, early diagnosis fungal nosocomial infections are often challenging due to lack of specific signs and late manifestation of symptoms which significantly render the efficiency of antifungal treaments^35^.

Considering these, although engineers and researchers are actively seeking for interventions that shape the indoor environment to be healthier for occupants, such efforts will not flourish until better undertesting is provided regarding interconnection between healthy building foundations. Among the nine foundations of healthy buildings, here we are particularly interested in the interaction of indoor light and indoor pathogens. Next to the psychological and physiological impact of daylight exposure on human body, light is an important environmental factor that shapes the microbial communities. As early as 500BCR, Egyptians have used the sunlight to treat chronic ulcers successfully^14^. Sunlight therapy has been in practice among communities worldwide to treat various diseases before the discovery of ultraviolet radiation (UV), which provides the foundation of today’s antimicrobial light therapy^14^. One of the breakthrough discoveries of photochemistry can be traced back to 1903 when Niels Ryberg won the Nobel Prize for using the Finsen lamp (>380nm) for treatments of skin tuberculosis^15,16^. The blue-light emitting lamp (400-500nm) effectively eradicates *Mycobacterium tuberculosis* and other pathogenic bacteria such as *P. aeruginosa, Methicillin-Resistance Staphylococcus aureus strains (MRSA), and A. baumanni*, which saved many lives subjected to potentially lethal burn during that time^15–19^. Although other spectrums of the visible light have important biological relevance to human, majority of the antimicrobial light studies focused on the violet-blue light spectrum (400-500nm) as it emits the highest energy within the visible light range. Blue light therapy has shown to be effective against a wide range of microorganisms including bacteria, fungi, virus, and yeast without the need of additional photosensitizers, which could be a promising intervention for disease controls in the future^14,19–23^. However, to date, it is not systematically well understood whether changing indoor light intensity and bringing more blue light into spaces can impact the viability of indoor microbiota on different surfaces.

To partially fulfill these gaps, using a controlled living-lab set-up, we investigated the effect of indoor daylight, modulated by electrochromic (EC) window, on the growth rate and viability of four important surface borne pathogenic bacteria *MRSA, P. aeruginosa, K. pneumoniae* and *E. coli*, as well as three indoor pathogenic fungal species *Stachybotrys chartarum, Aspergillus fumigatus*, and *Aspergillus versicolor* on different surface materials. Indoor light exposed groups were compared with window and blind conditions to evaluate the indoor light effects.

## 2 METHODS

### 2.1 Experimental chamber setup

All experiments were performed inside a controlled mini-living lab set-up. EC smart windows glass were installed on the chamber panels (Figure 1A). All glasses contain electrochromic coating, allowing window tints to be manually controlled. A custom solar simulator (Sciencetech) consisting of a xenon arc lamp and a series of optical filters used to illuminate the inside of the chamber with simulated sunlight with the light spectrum of natural sunlight (Figure 1A). Indoor daylight levels were controlled by EC window tints, with EC window-Clear (60% light transmission) and EC window-Tinted (1% light transmission) tested in this study. EC window-Tinted showed excellent glare control and had similar light intensity as the Blinds condition. Temperature and relative humidity inside the chamber were maintained within a narrow range for testing, specifically 24.7± 1 °C and 42 ± 3%, respectively.

**Figure 1:**
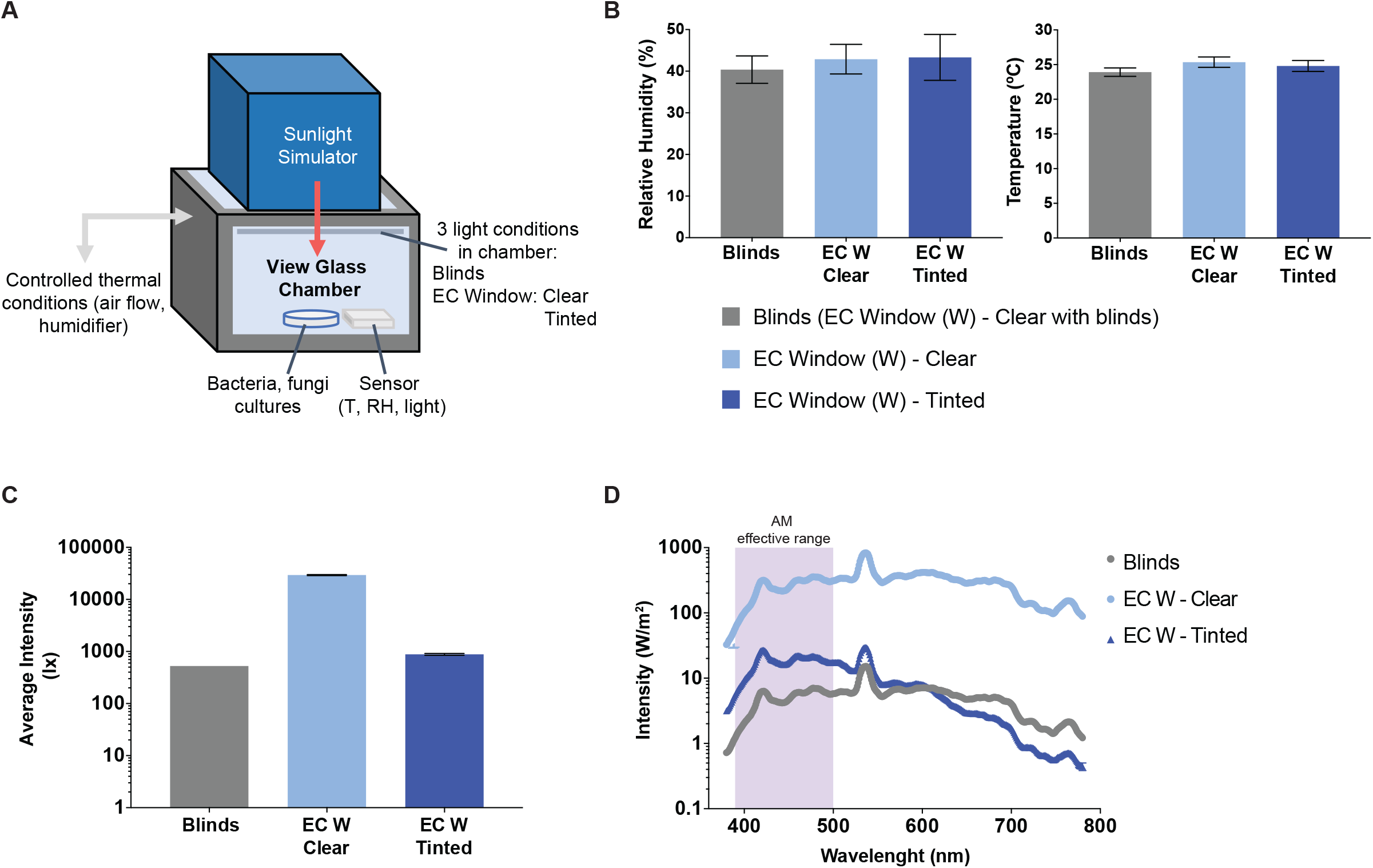
Experiment set up and conditions. **A)** Environmental chamber with EC Windows from View Inc. and temperature and humidity control. Samples are placed inside the chamber for indoor light exposure. **B)** Bar graph with standard deviation (SD) of temperature (left) and humidity (right) for each light condition: Blinds (grey), Clear (light blue), and Tinted (dark blue). **C)** Bar graph with standard deviation (SD) of light intensity of each light condition: Blinds (grey), Clear (light blue), and Tinted (dark blue). **D)** Light spectrum of three tested light conditions: Blind condition (grey line); EC Window - Clear (light blue line), and EC Window - Tinted (dark blue line). Violet-blue light (400 - 500nm) spectrum are highlighted with purple.

### 2.2 Strains and media preparation

#### Bacteria

The selected bacteria and fungi strains were purchased from the American Type Culture Collection (ATCC) as follows: *Methicillin-resistant staphylococcus aureus (MRSA; ATCC 6538), Pseudomonas aeruginosa (strain Boston 41501; ATCC 27853), Escherichia coli (ATCC 11229), and Klebsiella pneumoniae (ATCC 1352)*.

*MRSA* and *P. aeruginosa* were seeded in sterile Tryptic Soy Broth (TSB, BD Diagnostic) and *K. pneumonias and E. coli* were seeded in sterile Difco™ Nutrient Broth (Fisher Scientific) and incubated at 37°C degree overnight prior to each experiment. Bacterial cell density was adjusted to approximately 1×10^5^ cells per ml based on the optical density reading at 600nm (OD_600_).

#### Fungi

Fungal species tested in this study were *Aspergillus fumigatus (ATCC 1022), Aspergillus versicolor (ATCC 11730) and Stachybotrys chartarum (ATCC 201867)*. All species were grown on Potato Dextrose Agar (PDA) (Difco™ Fisher) at 25°C. Spores were extracted from agar plates by flooding method with 10ml of autoclaved distilled water several times and stored at 4°C. Spore density for each fungal species were enumerated by an automated cell counter (Countness3, ThermoFisher) and aqueous fungal suspension with a concentration of 1×10^4^ spores per ml was prepared.

### 2.3 Indoor daylight effect on the growth rate of indoor bacteria and fungi using high nutrient agar plates

This part was designed to mimic conditions where pathogens have access to high amount of nutrients (here we used agar) and they could actively grow. For bacteria, agar plates (n=3 per bacterial species per light condition) were inoculated with known bacterial concentration and placed inside the environmental chamber for indoor daylight exposure of 24 hours (T = 24.3°C, RH = 41%; Suppl Fig1A). Colonies forming units (CFU) on each plate were then counted. Negative controls (n=3 per bacterial species per light condition) were inoculated plates kept in the dark at 25°C (no indoor light exposure).

Similarly, PDA plates (n=3 for each fungal species) were inoculated with known concentration of fungal spores and exposed to the indoor daylight for 72 hours (T = 25.2°C, RH = 42; Suppl Fig1A). Area of fungi mycelium growth on each plate was then measured to assess the light effects. Negative controls (n=3 per fungal species per light condition) were inoculated plates kept in dark at 25°C (no indoor light exposure).

Both bacteria CFU and area of mycelium growth on each plate were processed through semi-automated image analysis (NIH, ImageJ) for quantitative assessment. Workflow can be found in Suppl Fig 1A, Suppl Fig 2A. Bacterial and fungal growth rate (CFU per ml or area of mycelium) at EC window (both clear and tinted) were compared with Blinds individually. Percentage of growth rate reduction was calculated with Equation 1. Results were summarized in Table 1.

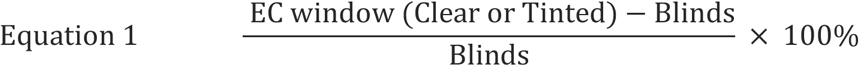

**Table 1.**
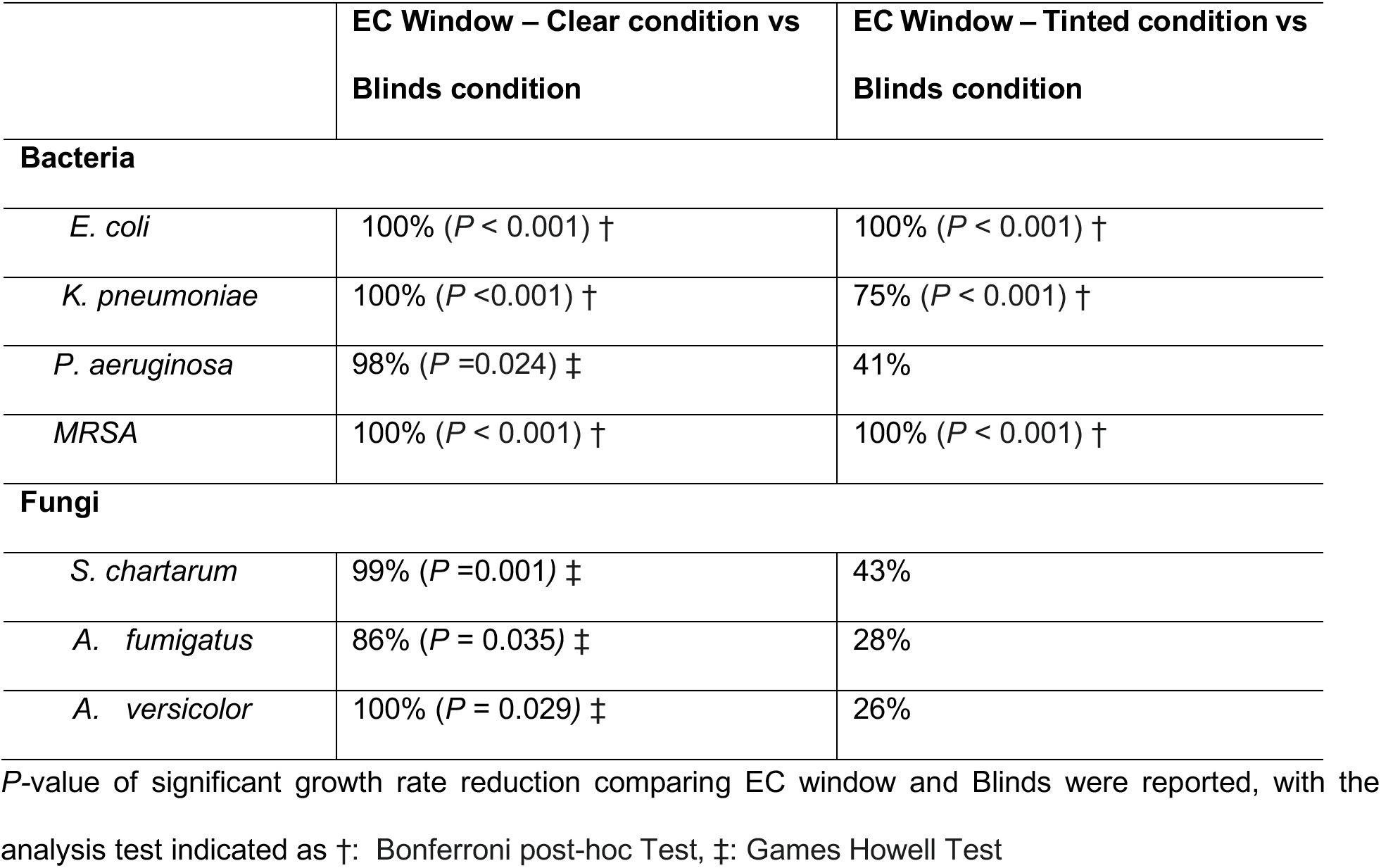
Percentage of reduction of bacterial CFU per ml and fungi mycelium growth by indoor daylight at EC Window – Clear and EC Window – Tinted on high nutrient agar plates.

### 2.4 Indoor daylight effect on the viability of indoor bacteria and fungi using low nutrient surface materials

This part was designed to mimic conditions where pathogens do not access to high amount of nutrients, and they could not actively grow. Specifically, the indoor daylight effect on bacteria and fungi on inanimate surfaces were assessed by inoculation of known concentration of species on selected indoor surface materials: polystyrene (PS), polyvinyl chloride (PVC) fabric, and glass. All materials were autoclaved and performed with triplicates for each tested species.

For bacterial, 2ml of the bacterial suspension (10^5^ cell/ml) were placed onto the sterile tested surface material (PS, PVC fabric, Glass) and placed inside the chamber for indoor daylight exposure of 24 hours (T = 24.3°C, RH = 42; Suppl Fig 1B). After exposure, bacteria from each sample were serially diluted and plated onto the agar plate (Tryptic Soy Agar for *MERS* and *P. aeruginosa*; Nutrient Agar for *E. coli and K. pneumoniae)*. The plates were then incubated at 37°C overnight and the CFU were counted. Negative controls (n=3 per fungal species per light condition) were inoculated plates kept in dark a 25°C (no indoor light exposure). Positive controls were inoculated agar plates kept at 37°C incubator.

For fungi, similar procedures were done with minor modifications: 3ml of aqueous fungal suspension with a concentration of 1×10^4^ spores/ml were placed on the three tested surface materials, placed inside the environmental chamber, and exposed to indoor daylight for 72 hours (T = 24.7°C, RH = 41%; Suppl Fig 1B). After exposure, samples were inoculated onto the PDA and incubated at 25°C.

After incubation, the bacteria CFU and area of fungal mycelium growth of each sample were quantified using ImageJ software. The workflow for nutrient poor experiment was included in the supplemental materials (Suppl Fig 1B, Suppl Fig 2A).

Bacteria and fungi viability reduction were also calculated for each material by comparing the EC window condition (Clear and Tinted) and Blinds. Percentage of reduction was calculated with Equation 1 and summarized in Table 2.

**Table 2.**
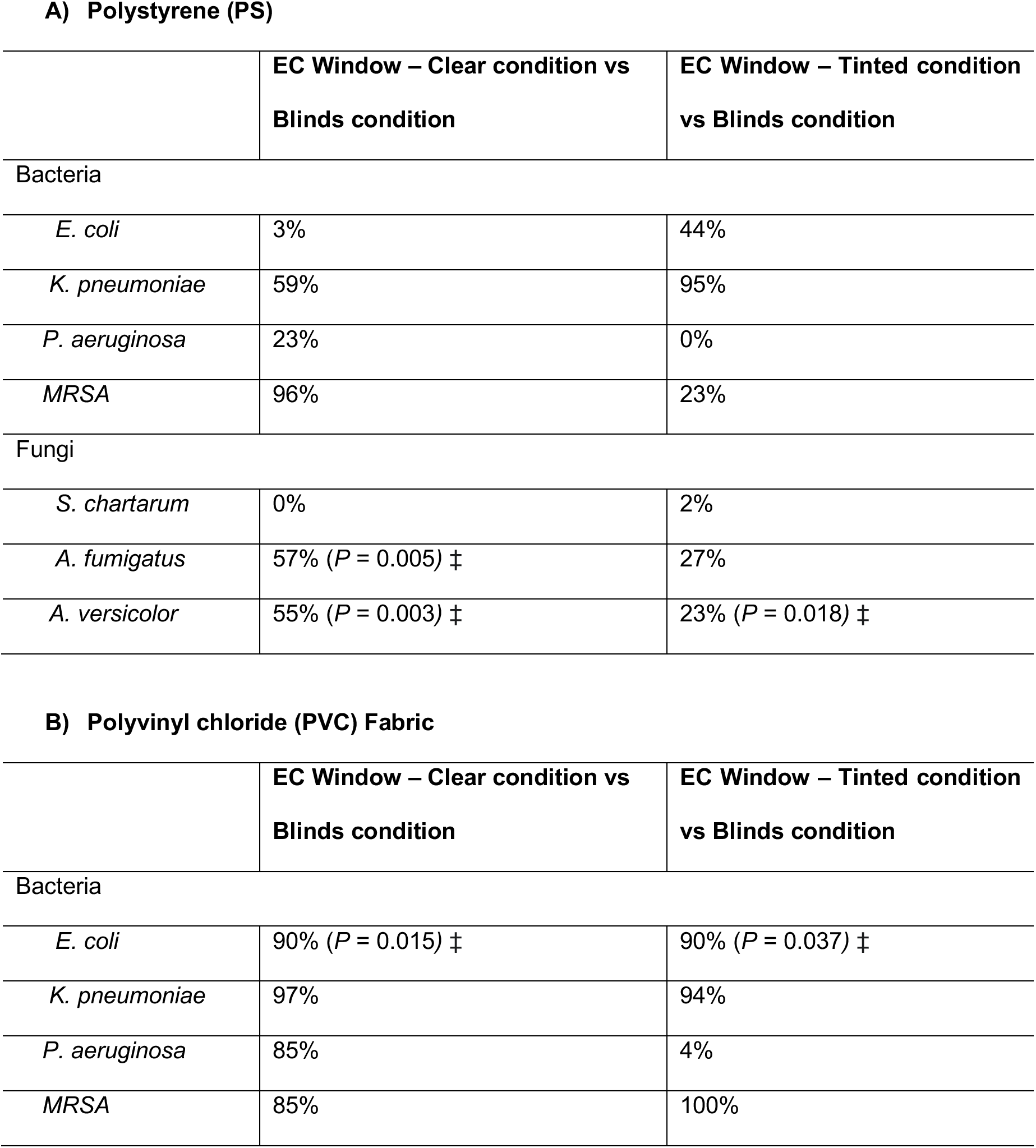
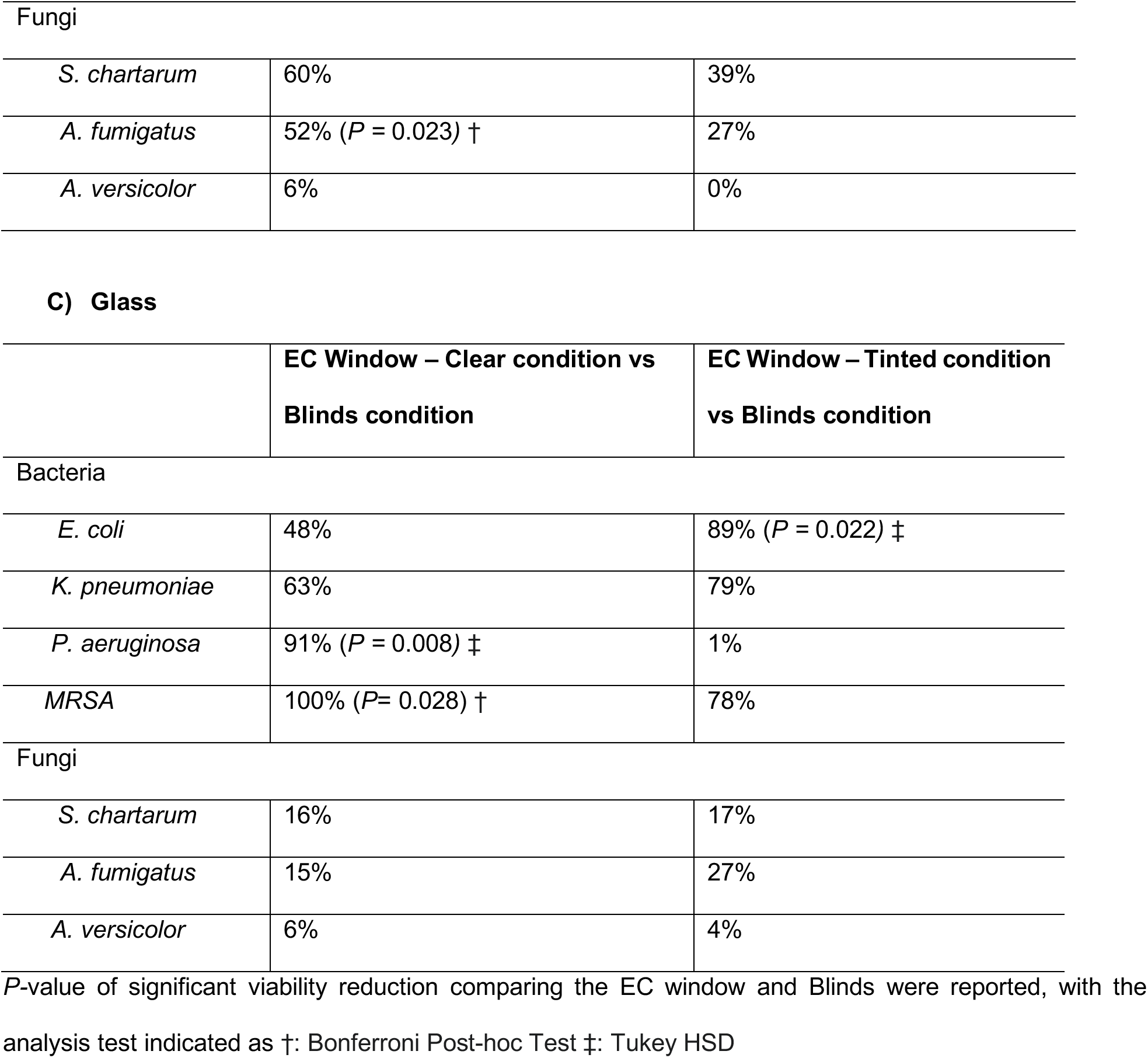
Percentage of reduction of bacterial CFU per ml and fungi mycelium growth by indoor daylight in EC Window – Clear condition and EC Window – Tinted condition relative to Blinds condition on A) Polystyrene B) PVC fabric and C) Glass.

**A) Polystyrene (PS)**
|  | EC Window – Clear condition vs<br>Blinds condition | EC Window – Tinted condition<br>vs Blinds condition |
| --- | --- | --- |
| Bacteria |  |  |
| *E. coli* | 3% | 44% |
| *K. pneumoniae* | 59% | 95% |
| *P. aeruginosa* | 23% | 0% |
| MRSA | 96% | 23% |
| Fungi |  |  |
| *S. chartarum* | 0% | 2% |
| *A. fumigatus* | 57% (*P* = 0.005) ‡ | 27% |
| *A. versicolor* | 55% (*P* = 0.003) ‡ | 23% (*P* = 0.018) ‡ |

**B) Polyvinyl chloride (PVC) Fabric**
|  | EC Window – Clear condition vs<br>Blinds condition | EC Window – Tinted condition<br>vs Blinds condition |
| --- | --- | --- |
| Bacteria |  |  |
| *E. coli* | 90% (*P* = 0.015) ‡ | 90% (*P* = 0.037) ‡ |
| *K. pneumoniae* | 97% | 94% |
| *P. aeruginosa* | 85% | 4% |
| MRSA | 85% | 100% |
| Fungi |  |  |
| *S. chartarum* | 60% | 39% |
| *A. fumigatus* | 52% ($P = 0.023$) † | 27% |
| *A. versicolor* | 6% | 0% |

C) Glass
|  | EC Window – Clear condition vs<br>Blinds condition | EC Window – Tinted condition<br>vs Blinds condition |
| --- | --- | --- |
| Bacteria |  |  |
| *E. coli* | 48% | 89% ($P = 0.022$) ‡ |
| *K. pneumoniae* | 63% | 79% |
| *P. aeruginosa* | 91% ($P = 0.008$) ‡ | 1% |
| MRSA | 100% ($P = 0.028$) † | 78% |
| Fungi |  |  |
| *S. chartarum* | 16% | 17% |
| *A. fumigatus* | 15% | 27% |
| *A. versicolor* | 6% | 4% |

$P$ -value of significant viability reduction comparing the EC window and Blinds were reported, with the analysis test indicated as †: Bonferroni Post-hoc Test ‡: Tukey HSD

### 2.5 Statistical analysis

All statistical analysis were conducted using statistical software base R 4.0.5 and all tests were given 5% risk and visualization were carried out using GraphPad Prism version 7.0a (Graph Pad Software, Inc, USA). Normality of data sets were assessed using the Shapiro-Wilk test. Homogeneity of variance was assessed using either Bartlette’s test or Levnene’s test depending on the normality.

For analysis of indoor light effect on bacteria and fungi growth rate on high nutrient agar plates, One-Way ANOVA followed by Bonferroni post-hoc test were used to compare between Blinds and EC windows (Clear and Tinted) since all our data followed Gaussian distribution (Shapiro-Wilk: *P*>0.05). Welch-corrected ANOVA was performed if equal variance assumption was not met (Bartlette’s test *P*>0.05), followed by Games Howell post-hoc test.

For analysis of indoor light effect on the viability of bacteria and fungi on low nutrient surfaces, Two-Way ANOVA was used to analyze the effect of light and material type if data was normally distributed (Shapiro-Wilk: *P*>0.05), followed by TukeyHSD test to compare Blinds with EC window condition. Kruskal Wallis test was used if data failed the normality test (Shapiro-Wilk: *P*>0.05), followed by Bonferroni post-hoc test. Results with *P*<0.05 were considered statistically significant and shown in Figure2, Table 1 and Table 2.

**Figure 2:**
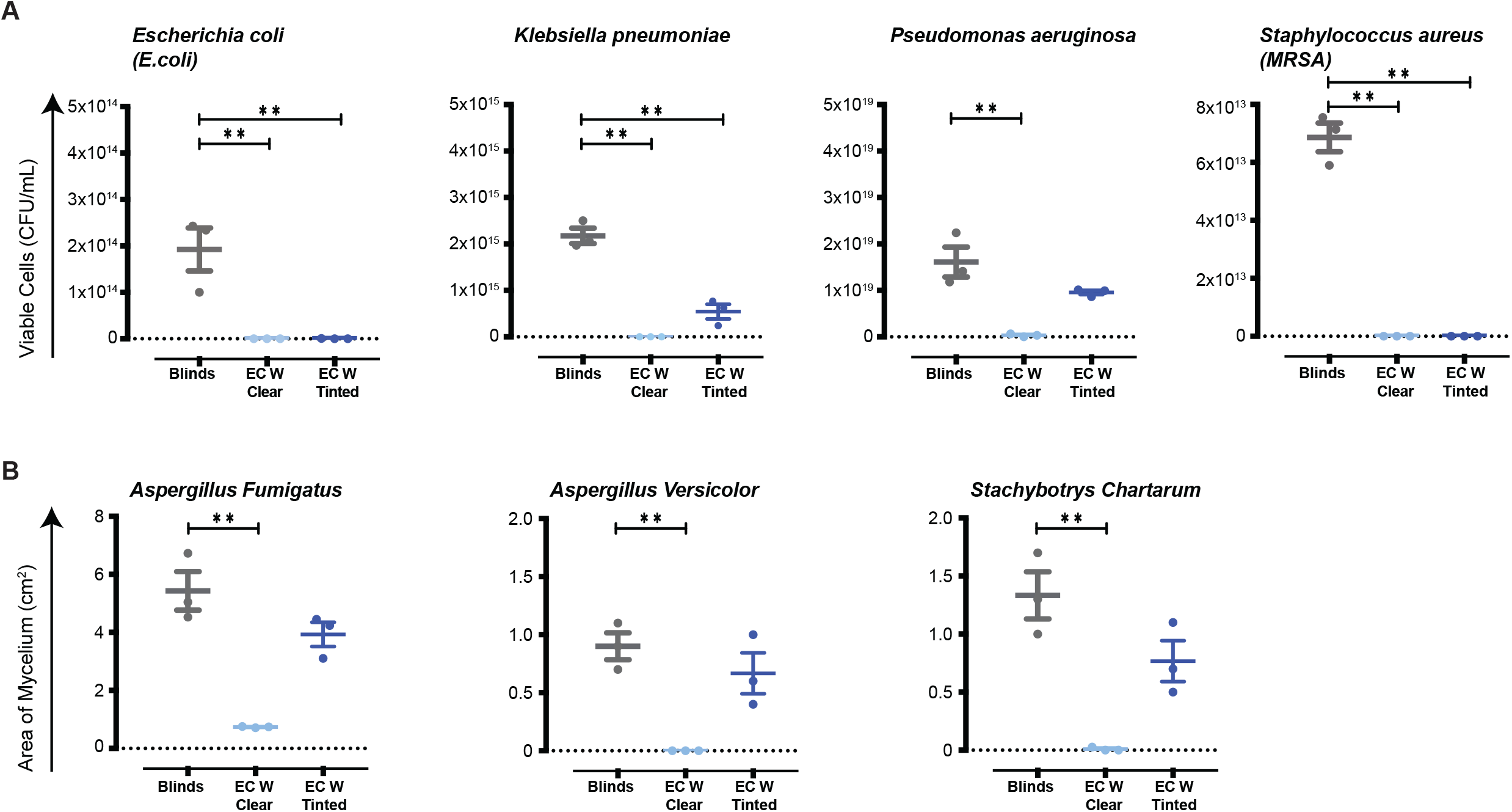
Bacteria and fungi viability on high nutrient agar plates after exposure of indoor light at different EC window conditions. **A)** Bacteria colonies forming units (CFU) per ml on high nutrient agar plates after 24 hours indoor daylight exposure. **B)** Fungi mycelium growth on high nutrient agar plates after 72 hours of indoor daylight exposure. Statistical analysis compared each condition relative to the Blind condition (grey) (*, *P*<0.05; **, *P*<0.01).

## 3 RESULTS

We developed a mini-living lab with electrochromic (EC) windows (Figure 1A); bacteria and fungi were placed on different surfaces and exposed to indoor daylight at different EC window tints inside the environmental chamber. For each experiment, the environmental chamber’s temperature and relative humidity (RH) were kept between 24.7°C ± 1°C and 42%± 3%, respectively (Figure 1B).

Indoor daylight was controlled by different EC window tints, corresponding to 60% light transmission (EC Window-Clear) and 1% light transmission (EC Window-Tinted). Light conditions at each window tint were shown in Figure 1B. The reference condition was 1% openness blackout roller shades that fully cover the top chamber glass in its clear state. Focusing on indoor daylight intensity, results showed that EC window at its clear state have an average light intensity of 29×10^3^ (lx), which was much higher than tinted EC window and Blinds (880 and 526 (lx), Figure 1C). In additions, while the tinted EC windows and blind conditions both let in 1% of light, the tinted windows preferentially let in shorter wavelength, higher energy daylight (400 - 500 nm; Figure 1D) while conditions with blinds let in higher wavelength, shorter energy daylight (550 - 650 nm; Figure 1D).

### 3.1 Indoor daylight effect on bacterial and fungal growth rate on high nutrient agar plates

To investigate the effect of indoor daylight, with different intensity and spectrum in response to window settings, on bacterial and fungal growth rate, microorganisms were cultured on high nutrient agar plates and exposed to different daylight conditions for 24 hours (bacteria) and 72 hours (fungi). Results showed that indoor daylight at EC window–Clear condition significantly reduced the viability and growth rate of all tested bacteria and fungi (One-way ANOVA; p-value < 0.05). Table 1 showed the percentage of reduction of bacteria CFU per ml and fungal area of mycelium by the two tested indoor daylight conditions compared to blinds. All tested bacteria viability were reduced more than 98% by indoor daylight in the EC Window – Clear condition and fungi mycelium growth were reduced more than 86% (Table 1) relative to blinds. The tinted condition significantly reduced the viability of *E. coli, K. pneumoniae* and *MRSA*, but not *P. aeruginosa*. The highest reduction was observed in *E. coli* and *MRSA*, in which the growth rate was reduced by 100% in both daylight conditions compared to blinds (Table1). Lowest reduction was observed in *P. aeruginosa*, which the growth rate was reduced by 41% at tinted condition.

For fungi, analysis showed significant reduction in fungal growth rate when exposed to high intensity of indoor light where windows were not tinted and blind was not present (EC Window-Clear). Under low indoor light intensity condition, results showed fungal growth rate reduction, but not statically significant when exposed to daylight through the tinted EC windows compared to blinds; 42.5% reduction for *S. chartarum*, 27% reduction for *A. fumigatus*, and 26% reduction for *A. versicolor* (Table 1, Figure 2B).

### 3.2 Indoor daylight effect on bacteria and fungi viability on low-nutrient surface materials

To examine the effect of daylight when bacteria and fungi were placed on different indoor surface materials, the tested microbes were placed on PS, PVC fabric, and glass for 24 hours (bacteria) and 72 hours (fungi).

The results showed that majority of bacterial and fungal species were highly viable when windows had blinds; while lower viability was mostly detected when EC Window were present (Figure 3), either viability reduced in response to higher indoor intensity under EC Window - Clear condition or presence of blue light spectrum under EC Window - Tinted condition. However, the extent of indoor light effect also depends on type of surface materials as well as type of species applied (factors interaction). For example, focusing on bacteria, *E. coli*, as a gram-negative bacteria, its growth rate was significantly reduced under both EC Window - Clear and EC Window-Tinted condition on PVC fabric (Turkey HSD p-value = 0.015 and p-value = 0.037, respectively). However, on glass surfaces, significant growth rate reduction was only observed under EC Window-Tinted condition (Turkey HSD; p-value = 0.022). No significant reduction was observed when *E.coli* was on PS (Table 2A). In contrast, viability of *MRSA*, as a gram-positive bacteria, was higher under blind condition, especially when they were on glass materials (EC Window-Clear: 100% reduction; Pairwise t-test with Bonferroni correction; p-value = 0.028, Table 2C). Interestingly, EC window conditions in its clear state reduced the viability of *MRSA* by 95.5% and 100% on PS and glass, respectively, and tinted condition reduced *MRSA* by 100% on PVC fabric (Table 2).

**Figure 3.**
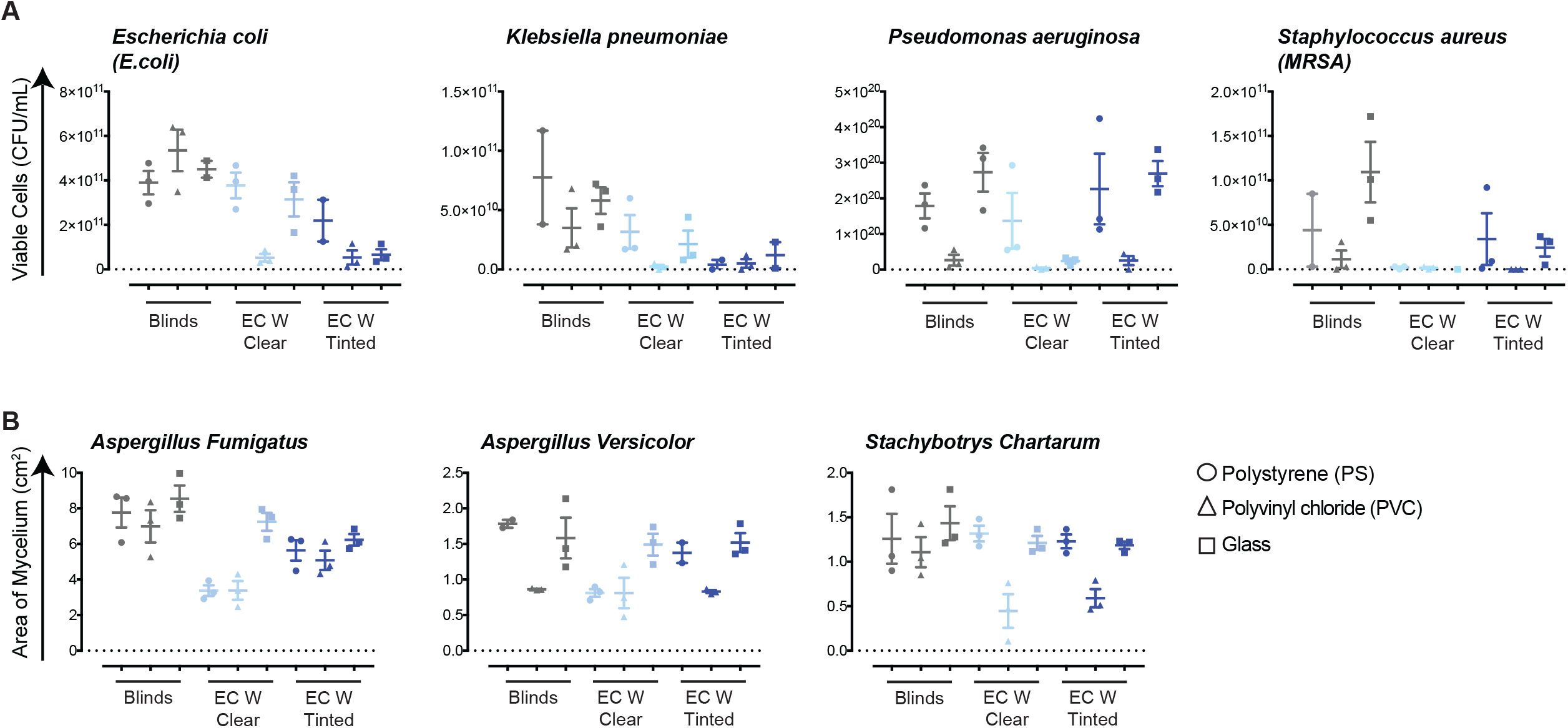
Effect of indoor daylight at different EC window conditions on bacteria and fungi variability on three different surface materials. **A)** Bacteria colonies forming (CFU) per ml on PS, PVC fabric and glass after 24 hours indoor daylight exposure. **B)** Fungi mycelium growth on PS, PVC fabric and glass after 72 hours indoor daylight exposure.

Viability of fungal pathogens was mostly reduced in response to higher indoor light intensity under EC - Clear condition, especially when they were on PS; 57 % reduction for *A. fumigatus* and 55 % reduction for *A. versicolor*, (Turkey HSD; p-value = 0.005 and 0.003 respectively (Figure 3 and Table 2A). EC Window–Tinted condition also reduced viability of *A. fumigatus* and *A. versicolor* on PS materials, but to a lesser degree (27% to 23%, respectively). In contrast, EC window (both Clear and Tinted state) reduced the viability of *S. chartarum* on PVC materials but did not reach a statical significance (Figure 3 and Table 2B); EC Window – Clear achieved a 60% viability reduction and EC Window – Tinted achieved 39% reduction (Two-Way ANOVA; p – value = 0.056).

## 4 DISCUSSION

Since indoor daylight and indoor microbiomes both have important relevance to occupant health, it is worth investigating their interactions. Our work indicates that indoor daylight modulated by the EC smart window can significantly reduce the viability and growth rate of pathogenic bacteria and fungi, especially when nutrients are available for active growth, mainly due to exposure to shorter wavelength and higher energy spectrum indoor light for bacterial pathogens. Interestingly, fungi mycelium growth and viability were more affected by light intensities rather than light spectrum, where light effect were mostly seen at EC-window-Clear condition when highest daylight intensities were transmitted.

Specifically, we found that the growth rate of all tested pathogenic bacteria on high nutrient condition were significantly reduced by daylight in both EC Window-Clear and EC Window-Tinted conditions. The results were expected since significant amount of high energy blue light (400-500nm) were transmitted through EC window tints relative to blinds, despite the tinted condition and blinds both having 1% overall light transmission. The bactericidal effect of blue light at 400-500nm have been extensively reported in many papers^21,23,39–43^. Bacteria employs different types of photosensitizers for blue light sensing, with some commonly found examples being flavins, porphyrins and NADH^21^. The absorption efficiency of cytochromes directly determines the effectiveness of antimicrobial blue light, which various between each species at the range of 400-500nm. Exposure of blue light at this range excites the cytochromes and induces a cascade of oxygen dependent photoexcitation reactions^40,41^. The generated Reactive Oxygen Species (ROS) can cause substantial oxidative damages, disrupt cellular functions and lead to cell death if accumulates in cells^16,20,23,39,41^.

Likewise, reduction of fungi mycelium growth by indoor daylight in the EC Window-Clear condition was also expected as fungi light response has long been understood^44,45^. Blue light response has been well-described in several *Aspergillus* species, which agreed with our finding that continuous white light and blue light exposure inhibits conidia germination and fungi mycelium growth^22,46,47^. However, daylight in the EC Window-Tinted condition did not significantly reduce the fungal mycelium growth. Unlike bacteria that uses cytochromes, blue light sensing in fungi involves the white-collar complex (WCC) which is not affected by its absorption efficiencies^48^. Secondly, fungi are known to produce pigments as first line of defense to photoinactivation^49,50^. Melanin is one of the photoprotective pigments that are found in many fungi species. The hydrophobic pigment serves as a scavenger for ROS and reactive nitrogen species (RNS) to protect the cells, which are important for UV and visible light protection in fungi^49,51^. 1,8-dihydroxynaphthalene (DHN) melanin are found in the conidia of *Aspergillus*, which is responsible for the gray-greenish color of the mycelium^50^. Multiple light defense mechanisms make fungi more resistant to indoor light compared to bacteria, thus higher doses of antimicrobial blue light (400-500nm) are needed for reducing fungal mycelium growth. As daylight in the EC-Tinted condition shows much lower blue light intensity than the EC Window-Clear condition, it is not surprising that daylight in this condition (EC-Tinted) did not have a significantly potent effect on fungi growth. Studies have found that the impact of blue light is often at much higher light intensities^22,46,52^. Hatakeyama et.al (2017) exposed *Aspergillus oryzae* culture on high nutrient agar plates to continuous blue light at 430nm, and found that the fungal mycelium growth and number of fungal conidia were significantly decreased compared to those grew in the dark^52^. Note that the blue light intensity used in their study (94μmol m^−2^ s^−2^ for both) is much higher than our study (5W/ m^−2^ for the EC Window-Tinted condition). Nevertheless, light still plays a critical role in regulating fungal biological activities, including conidial germination rate, sexual development and circadian rhythm, and thus should be taken into consideration for fungal disease transmission^45,46,48,53^.

We also investigated the effect of indoor daylight on the viability of bacteria and fungi under low nutrient condition using different surfaces materials: polystyrene (PS), polyvinyl chloride (PVC) fabric and glass. Our results showed that bacteria and fungi microbial viability were significantly lower on PVC fabric compared to glass and PS. This could be explained by the materials properties, as porous material (PVC fabric) entraps microorganisms within the matrix hence less viability bacteria were recovered compared to those on non-porous surfaces^30,54^. Another explanation of the low bacteria viability on PVC fabric could be the presence of chemical dye. Industrial textile often employed chemical agents and coatings that have known antimicrobial properties, which targets a wide range of microorganisms including bacteria, fungi and virus^55^. These type of fabrics are frequently used in high traffic built environments such as hotels and hospitals, where materials such as towels, curtains, and carpets could all potentially serve as fomites that facilitate disease transmission^55^. However, we are not aware of any antibacterial agents on the PVC blind material tested in this study.

Our study also found that gram-positive bacteria (*MRSA*) were more susceptible to indoor daylight than gram-negative bacteria. As mentioned in the previous section, the type of porphyrins employed by the bacteria for blue-light sensing determine the photoinactivation efficiency, which varies at the strain levels^23^. The predominant porphyrin produced in *S. aureus* is coproporphyrin. In contrast, various types of porphyrins are found in gram-negative bacteria^23^. This could also explain the variability in blue light effectiveness between the bacterial species. Several studies have also reported the higher blue light susceptibility of gram-positive bacteria than gram-negative bacteria^20,23^, though contradictory findings were also reported: Guffey & Wilborn (2007) found that gram-positive bacteria (*S. aureus*) were more resistant to blue light inhibition than gram-negative bacteria (*P. aeruginosa*), which could be due to different experimental designs (starting concentration, exposure time, blue light intensity) and different bacterial strains used in the study^56^. Further investigations are needed to explain the contradictory findings between studies.

For fungi viability testing on different surface materials, significant fungal viability reductions were only observed on *A. fumigatus* on PS in both EC Window conditions. No light effects were found on glass for any of the three tested fungi species. Although fungi blue light studies are relatively sparse compared to bacteria, some assumptions can be made based on the characteristics of the fungi: Fungi in genus *Aspergillus* contains filamentous hyphae and conidia that are highly hydrophobic^51,57^. The shape and properties of fungal spores various between species, and the chemistry and physiochemistry of surface materials also influence conidial binding ability hence affecting the indoor daylight effect^58^. We also did not detect any significant indoor daylight effects on *S. chartarum*, which is in line with previous findings that *S. chartarum* has high light-resistance. A study by Green et.al (2005) found that ultraviolet germicidal irradiation (UVGI) at 265nm (144mJ cm^−2^) was not efficient to inactivate 90% of the *S. chartarum* spores^59^. It should note that UVGI emits the highest energy among the UV spectrum, which is much stronger than visible blue light^59^. Therefore, it is not surprising that indoor daylight used in this study was not efficient in reducing the viability of *S. chartarum*.

Overall, this study provides important insights for fomite transmission and healthy building design for several reasons. Extensive studies have shown that pathogenic bacteria and fungi can persist on inanimate surfaces for prolonged period of time, leading to disease transmission^30,60–62^. Especially in healthcare settings, numerous healthcare-associated infections and outbreaks have been associated with patient’s care items, ranging from personal items such as computer keyboard and tablets, to common high touch surfaces such as curtain, window, hand sanitizers dispensers, and medical devices such as, medical chart and thermometers^33^. The three tested materials (plastic for PS, PVC fabric for textile, and glass) in this study are frequently used in building materials, furnishings and devices, which could serve as a reservoir for indoor pathogen that causes nosocomial infections^63^. Moreover, the selected bacteria and fungi are all pathogenic species that are known to cause nosocomial infections, with the exception of *S. chartarum* which are known to produce various mycotoxins and often related to Sick Building Syndrome^64^. Additionally, among the tested bacteria pathogens, three (*MRSA, P. aeruginosa, K. pneumoniae*) of them belongs to EKSAPE pathogens which are biofilm-forming multidrug resistant organisms. We tested more gram-negative bacteria as majority of the HAI outbreaks are associated with gram-negative rods bacteria such as the *P. aeruginosa, K. pneumoniae* and *E. coli*, which are what we tested in this study^33^. This highlights the importance of our results as disinfections of antibiotic resistant pathogens are becoming increasingly challenging, considering nearly every current antibiotics have been observed with microbial resistance^35^. In addition to the potential influence on HAIs, daylight and access to views can shorten hospital length of stay, reduce pain medication use and improve overall patient experience, resulting in higher patient satisfaction ratings^65,66^. Patients are 47% more likely to choose a hospital room that has EC Windows than one that has blinds.

However, it should note that hospitals are only one of the many types of built environments in the society. Fomite transmission and other indoor pathogens also presents in all indoor environments such as offices, schools, and homes where people spend significant amount of their time. Currently, many building elements are still far from optimized, with regular commercial windows being an example that transmits excessive glare and heats. EC-window technology shows promising potentials to replace blind as the predominant glare control strategy, which previous studies have shown to improve cognitive function and psychological health of the occupants^4,7,13^. This study further expands its potential in minimizing disease transmission, which could be implemented in various built environments for shaping healthy indoor microbiomes towards the occupants.

### 4.1 Strength & Limitations

There are several notable strengths in our study design. First of all, this is the first study that investigated the interactions between indoor daylight and indoor microbiomes on different surfaces. Secondly, we utilized a highly controlled laboratory environment to simulate real indoor environments while still limiting the potential for confounding by temperature, humidity or ventilation. Current understandings of pathogenic persistence on inanimate surfaces is limited hence this study provides valuable insights on disease transmission. However, several limitations are also present. Fungi culture was only incubated for 72 hours (3 days), which may not be efficient for fungal mycelium to be fully developed. Longer exposure and incubation periods may be needed to see the indoor daylight effects. Furthermore, microbial contaminations are often caused by pathogens that dry on inanimate surfaces, where this study uses liquid bacterial and fungal spore culture for viability assessments due to the recovery limitations. It would also be valuable to test other surface materials such as stainless steel and textiles (clothes) to expand the understandings of indoor daylights effects on other type of high touch surface materials. Additionally, more gram-positive pathogenic bacteria can be tested as we only tested *MRSA* in this study. Finally, this study simulates a real world setting and does not necessarily reflect the germicidal effect that would be found in practice. The study focuses on the viability of bacteria and fungi rather than the risk of transmission or colonization. Research extending these findings to actual buildings or developing epidemiological models to estimate infection risk could extend the implications of these findings. Nevertheless, this study filled multiple knowledge gaps and provided important insights for future healthy building research.

## 5 CONCLUSION

In summary, we found that daylight passing through electrochromic (EC) windows, both in their clear state and tinted state, resulted in significant disinfection of bacteria on high nutrient surfaces relative to daylight passing through a clear window with blinds. This research shows that antimicrobial daylight in the 400-500nm range of the visible spectrum can limit the viability of bacteria and growth of fungi. Bacteria were highly sensitive to the light wavelength with significantly less viability when exposed to shorter wavelength light through a tinted window than the same intensity of light through a blind. Light intensity had stronger effects on fungi than light spectrum, which fungi mycelium growth and viability were only reduced at EC-Clear conditions. Indoor daylight effects various depends on the material types and microorganisms. Gram-positive bacteria (*MRSA*) were found to be more susceptible to indoor daylight compared to gram-negative bacteria, due to the different types of blue light photosensitizer employed by the species. Bacteria viabilities were significantly lower on porous material (PVC fabric) compared to non-porous materials (Glass & PS), which could be because of the material properties or chemical agents used in the textile. This study filled multiple knowledge gaps and showed the potential of EC window as an important technology to mitigate pathogen viability in office, residential, aviation and healthcare settings.

## ACKNOWLEDGEMENT

This research was funded by MITACS grant IT21657. We acknowledge View Inc. for providing the EC Window chamber used in the experiments.

## Supplemental Materials

**Supplemental Figure 1:** Experimental workflow of **A)** Indoor daylight effect on bacteria and fungi growth rate on high nutrient agar plates. **B**) Indoor daylight effect of bacteria and fungi viability on low nutrient surface materials experiment.

**Supplemental Figure 2**: Bacteria viability assessment **A)** Bacteria colonies forming unit counting using ImageJ software. **B)** Images of bacteria viability at different daylight conditions and control settings. From left to the right: bacteria growth at: 37°C in the dark incubator, 25°C in the dark incubator; environmental chamber at window with blind condition, environmental chamber at EC window-Clear condition at 25°C, environmental chamber EC window-Tinted at 25°C.

**Supplemental Figure 3** Images of fungal mycelium growth viability at different light condition and control settings. From left to the right: bacteria growth at: 37°C in the dark incubator, 25°C in the dark incubator; environmental chamber at window with blind condition, environmental chamber at EC window-Clear condition at 25°C, environmental chamber EC window-Tinted at 25°C.

