## Supplementary figures and images for "The effect of indoor daylight spectrum and intensity on viability of indoor pathogens on different surface materials"

### Supplemental Figures

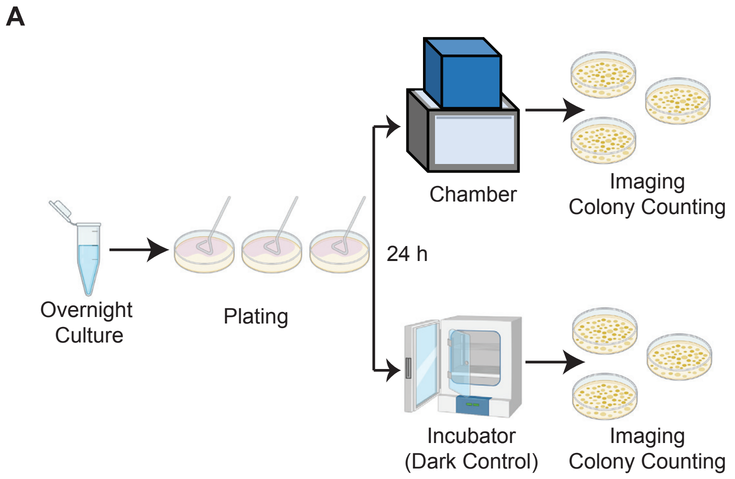

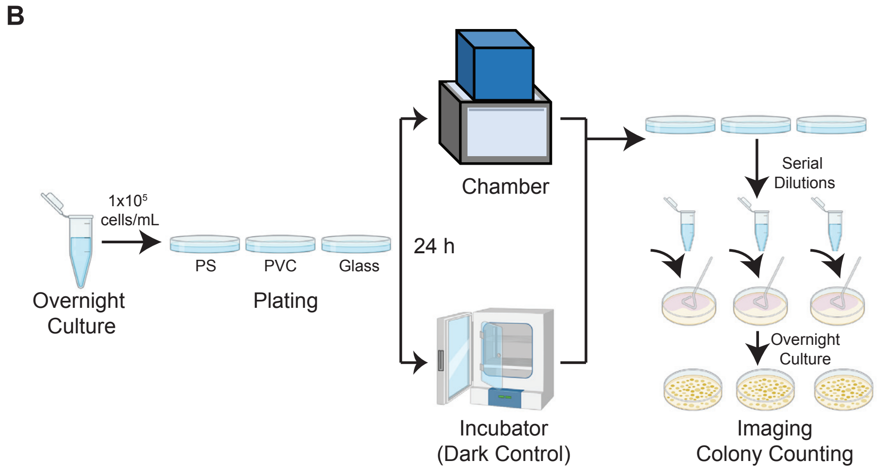

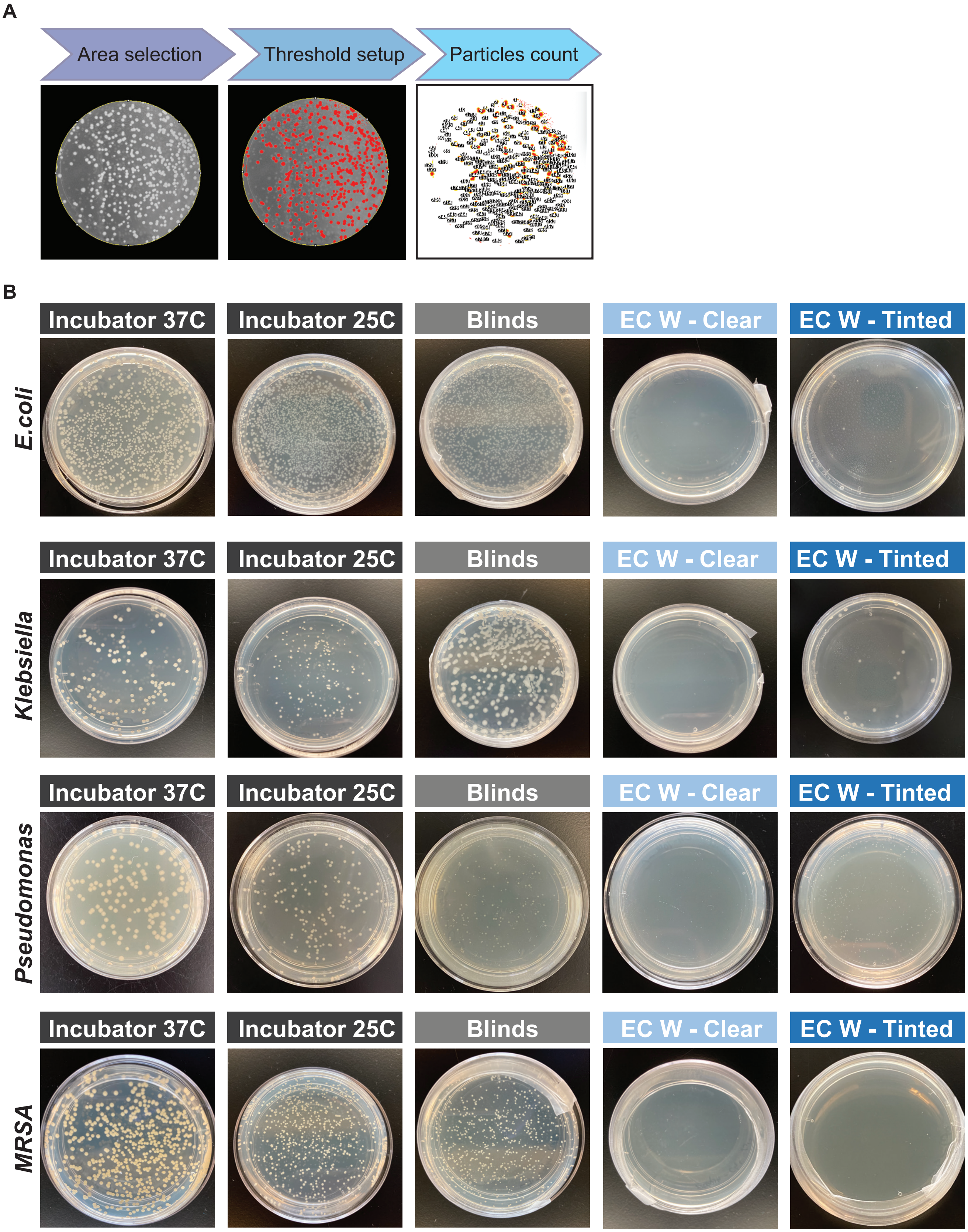

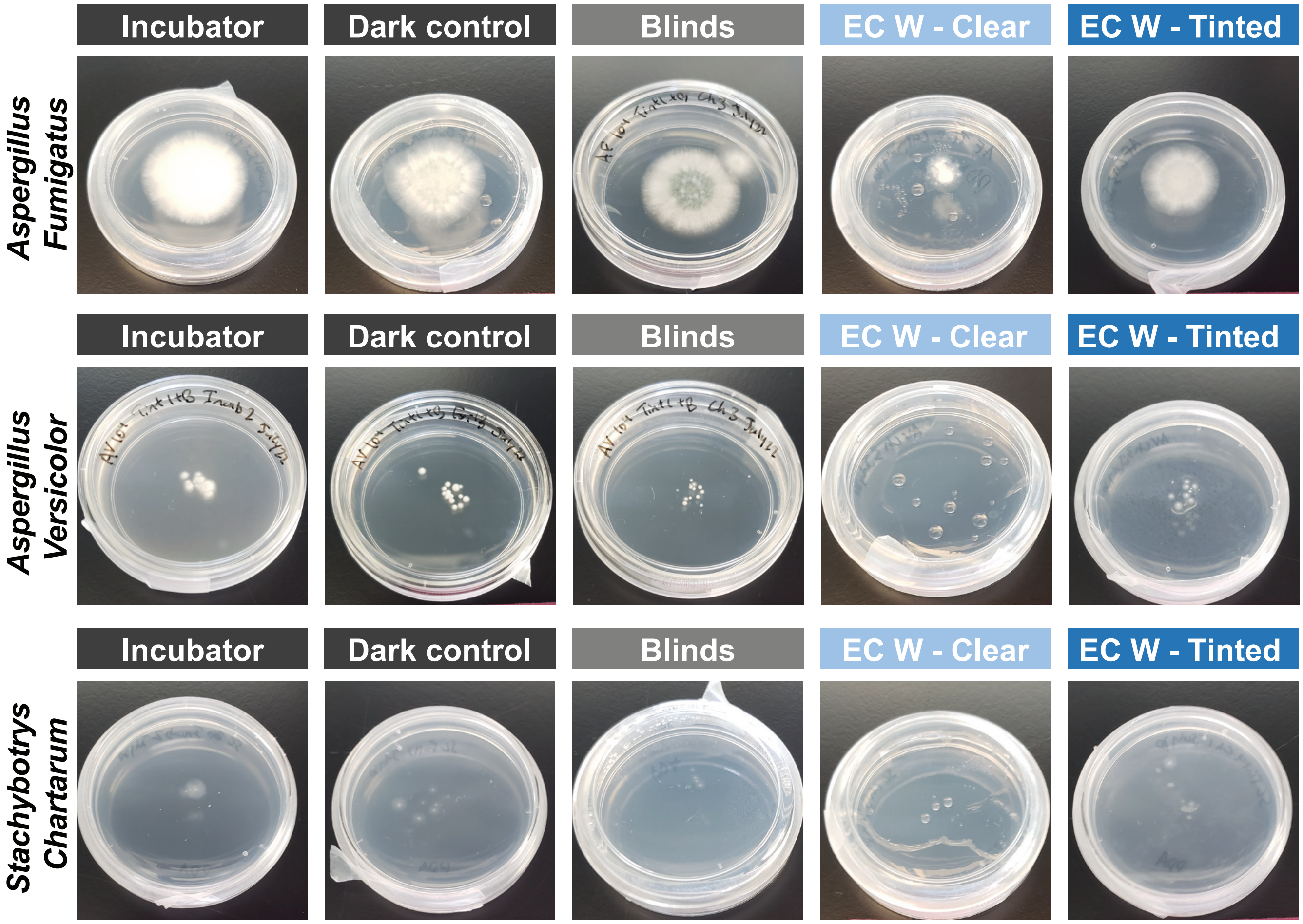
